# A map of *cis*-regulatory modules and constituent transcription factor binding sites in 80% of the mouse genome

**DOI:** 10.1101/2022.05.30.494043

**Authors:** Pengyu Ni, David Wilson, Zhengchang Su

## Abstract

Mouse is probably the most important model organism to study mammal biology and human diseases. A better understanding of the mouse genome will help understand the human genome, biology and diseases. However, despite the recent progress, characterization of the regulatory sequences in the mouse genome is still far from complete, limiting its use to understand the regulatory sequences in the human genome. Here, by integrating binding peaks in 9,060 transcription factor (TF) ChIP-seq datasets that cover 79.9% of the mouse mappable genome using an efficient pipeline, we were able to partition these binding peak-covered genome regions into a *cis*-regulatory module (CRM) candidate (CRMC) set and a non-CRMC sets. The CRMCs contain 912,197 putative CRMs and 38,554,729 TF binding sites (TFBSs) islands, covering 55.5% and 24.4% of the mappable genome, respectively. The CRMCs tend to be under strongly evolutionary constraints, indicating that they are likely *cis*-regulatory; while the non-CRMCs are largely selectively neutral, indicating that they are unlikely *cis*-regulatory. Based on evolutionary profiles of the genome positions, we further estimated that 63.8% and 27.4% of the mouse genome might code for CRMs and TFBSs, respectively. Validation using experimental data suggests that at least most of the CRMCs are authentic. Thus, this unprecedentedly comprehensive map of CRMs and TFBSs can be a good resource to guide experimental studies of regulatory genomes in mice and humans.

## INTRODUCTION

Mouse is probably the most widely used model organism to understand mammal biology and pathology of human diseases. Thus, it is not surprising that mouse is the first sequenced non-human mammal (Waterston et al. 2002). Conserved syntenies between the human and mouse genomes provide a powerful tool to understand functions of the human genome based on the known functions of the mouse orthologous sequences (Waterston et al. 2002). This homology-based approach plays a critical role in annotating the coding DNA sequences (CDSs) in the human genome. However, the power of such comparative genomics approach is hampered in annotating human *cis*-regulatory sequences due to the lack of a good understanding of mouse orthologous sequences and their less conservation nature compared with CDSs (Pennacchio and Visel 2010). c*is*-regulatory sequences such as promoters, enhancers and silencers are also called *cis*-regulatory modules (CRMs). While promoters are located upstream of target genes, enhancers and silencers can be far away (up to millions base pairs) from target genes, and they regulate the transcription of target genes independently of their locations and orientation (Levine and Tjian 2003; Davidson 2006). A CRM is made of clusters of transcriptional factor (TF) binding sites (TFBSs) of the same and different cooperative TFs, with a length ranging from hundreds to thousands of base pairs (Spitz and Furlong 2012). A CRM carries out its transcriptional regulatory function through specific bindings of cognate TFs to the TFBSs that it harbors. CRMs play equally important roles as CDSs in development, homeostasis, responses to environmental changes and evolution of organisms (Wray 2007). Diversity of CRMs may play even more important roles in driving diverse complex traits in humans (Siepel and Arbiza 2014) and mice (Attanasio et al. 2013). For example, genome-wide association studies (GWAS) in humans have found that most (90%) complex trait-associated single nucleotide variations (SNVs) reside in non-coding sequences (Hindorff et al. 2009; Ramos et al. 2014). Many of these SNVs overlap and disrupt TFBSs (Maurano et al. 2012), thereby affecting gene transcription (Kasowski et al. 2013; Kilpinen et al. 2013; McVicker et al. 2013; Wu et al. 2013; Huang and Ovcharenko 2015), and ultimately complex traits and diseases. On the other hand, many SNVs are in linkage disequilibrium (LD) with nearby TFBSs, and thus may not necessarily causal (Majewski and Pastinen 2011; Fu et al. 2013; Spielmann and Klopocki 2013; Herz et al. 2014; Ongen et al. 2014; Smith and Shilatifard 2014; Mathelier et al. 2015; Heyn et al. 2016; Khurana et al. 2016; Zhou and Wei 2016; Li et al. 2018). Hence, a better understanding of the mouse CRMs will not only aid to understand various aspects of mouse biology and make it an even better model of human diseases, but also will facilitate annotating human CRMs and understanding human biology. For example, studies of CRMs in human cells or tissues can be complemented by manipulating the orthologous CRMs in transgenic mouse in vivo using knockout and knockin techniques (Waterston et al. 2002; Visel et al. 2008; Visel et al. 2013).

In fact, great efforts have been made in the last decade to systematically annotate CRMs and constituent TFBSs in the mouse genome by the research community including the mouse ENCODE consortium (Shen et al. 2012; Yue et al. 2014; Moore et al. 2020) and individual groups worldwide using state-of-the-art techniques. Particularly, an enormous amount of data have been generated using ChIP-seq techniques to locate CRM function-related epigenetic marks (Boyle et al. 2008; Buenrostro et al. 2013) and TF bindings (Johnson et al. 2007) in the genomes of various mouse cell/tissue types. Numerous machine-learning methods (Ernst and Kellis 2010; Firpi et al. 2010; Hoffman et al. 2012; Rajagopal et al. 2013; Kleftogiannis et al. 2015; Gao et al. 2016; Gao and Qian 2020) have been developed to simultaneously predict CRMs and their functional states using location data of multiple epigenetic marks including histone modifications such as H3K3me1 (Dorighi et al. 2017; Rickels et al. 2017; Rada-Iglesias 2018), H3H4me3 (Howe et al. 2017) and H3K27ac (Zhang et al. 2020), and chromatin accessibility (CA) measured by DNase I hypersensitivity (Boyle et al. 2008) and transposase accessibility (Buenrostro et al. 2013). Although conceptually attractive, these methods suffer quite high false discovery rates (FDRs) (Kheradpour et al. 2013; Kwasnieski et al. 2014; Dogan et al. 2015; Kleftogiannis et al. 2015; Catarino and Stark 2018; Arbel et al. 2019; Ni and Su 2021) duo probably to low the specificity of these epigenetic marks used (Dogan et al. 2015; Young et al. 2017; Catarino and Stark 2018; Arbel et al. 2019; Ni and Su 2021) and at the same time, they might miss many CRMs in the genome because these data are only available in a few cell types (Ni and Su 2021). Moreover, these methods do not predict constituent TFBSs in the CRMs, notwithstanding it is mainly the TFBSs in a CRM that determine its functions (Spitz and Furlong 2012; Erceg et al. 2014). More recently, the ENCODE phase 3 consortium (Moore et al. 2020) predicted 339,815 candidate *cis*-regulatory elements (cCREs) in the mouse genome based on overlaps between millions of DNase I hypersensitivity sites (Thurman et al. 2012), transposase accessible sites (Buenrostro et al. 2013), active promoter histone mark H3K4me3 (Aday et al. 2011) peaks, active enhancer mark H3K27ac (Creyghton et al. 2010) peaks, and insulator mark CTCT(Kim et al. 2007) peaks, in a large number of mouse cell/tissue types. Nonetheless, the cCREs with an almost uniform length of 272bp are likely only fragments of full-length CRMs, because the known mouse enhancers have a mean length about 2,400bp (Visel et al. 2007). Moreover, the cCREs make up of 3.4% of the mouse genome (Moore et al. 2020), they might be largely under predicted.

To overcome the limitations of the existing methods, we proposed a two-step approach to first predict a map of CRMs and their constituent TFBSs in the genome using all available TF ChIP-seq data in the organism, and then predict functional states of all the predicted CRMs in any cell/tissue type of the organism using few epigenetic marks from the very cell/tissue type (Ni and Su 2021). We recently developed a new CRM predictor dePCRM2 (Ni and Su 2021) for the first step of our approach. dePCRM2 works by identifying closely located clusters of TFBSs in a genome through integrating all available thousands of TF ChIP-seq datasets in the organism (Ni and Su 2021). Unlike the existing methods, we use TF ChIP-seq data instead of CA and histone modification data to predict the loci of CRMs and constituent TFBSs, because it has been shown that TF binding is a more reliable predictor of CRM loci than CA and histone marks (Dogan et al. 2015). Using dePCRM2, we have predicted a highly accurate and unprecedentedly complete map of CRMs and constituent TFBSs in the human genome using then available 6,092 TF ChIP-seq datasets in hundreds of human cell/tissue types (Ni and Su 2021). In this study, we applied dePCRM2 to 9,060 TF ChIP-seq datasets for 701 TF in 438 mouse cell/tissue types, and predicted an unprecedentedly complete map of CRMs and constituent TFBSs in 79.9% of the mouse genome. Validation of the map using orthogonal evolutionary and experimental data suggests that our predictions are highly accurate. The map can be a good resource to guide experimental studies of the regulatory genomes of both mice and humans.

## RESULTS

### The 1,000pb binding peaks for cooperative TFs in different datasets have extensive overlaps

After filtering out low-quality peaks in the 9,060 TF ChIP-seq datasets (Table S1) collected (Materials and Methods), we ended up with 9,055 non-empty datasets for 701 TFs in 438 cell line/tissue/organ types. As in the case in humans (Niu et al. 2018; Ni and Su 2021), these available datasets are high biased to a few well-studied cell/tissue types (Figure 1A) and TFs (Figure 1B). For example, 1,020, 504 and 545 datasets were collected from mouse embryonic stem cells, epithelial cells and macrophage in bone marrow, respectively, while only one dataset was generated from 68 cell/tissue types, including pancreas beta cell MIN6B1, superior cervical ganglion, hepatocellular carcinoma, and so on (Table S1). Moreover, 460 and 160 datasets were collected for TFs Ctcf and SpI1, respectively, while just one dataset was produced for 131 TFs, such as Tfcp2, Nelfb, Hoxd11, and so on (Table S1). The number of called binding peaks in a dataset vary widely, ranging from 1 to 110,347, with an average of 15,069 peaks (Figure 1C). For instance, there are only 1, 4, 5 peaks for Top2B, Exh2, and Tbr1 in neuron, brain, and brain cortex, respectively, while there are 98,520, 99,344, and 110,347 peaks for Smc1A, Bcl6 and SpI1 in 3T3-L1 Adipocyte, helper cells and myeloid cells, respectively. The length of the called binding peaks also vary widely, ranging from 21 to 11,047 bp with a mean of 315bp, but the vast majority of them (98.7%) are shorter than 1,000bp (Figure 1D). We extracted 1,000bp genomic sequences centered on the summits of the called binding peaks for motif-finding to identify motifs of both the ChIP-ed TFs and their cooperators in each dataset (Li et al. 2019; Ni and Su 2021). Therefore, we extended the lengths of most (98.7%) of the originally called binding peak. We did so, because almost all known mammal enhancers with a mean length about 2,400bp (Visel et al. 2007) were longer than the mean length (315pb) of the called binding peaks and TFBSs are scattered along the entire lengths of enhancers (Zhang et al. 2011; Bailey and Machanick 2012; Sun et al. 2012; Li et al. 2019). We have previously shown that such extension (~1,000bp) of called peaks does not affect finding the motifs of ChIP-ed TFs, which typically reside in the middle of the peaks, but allows to find motifs of cooperative TFs, which can reside anywhere along the extended peaks (Li et al. 2019; Ni and Su 2021).

**Figure 1.**
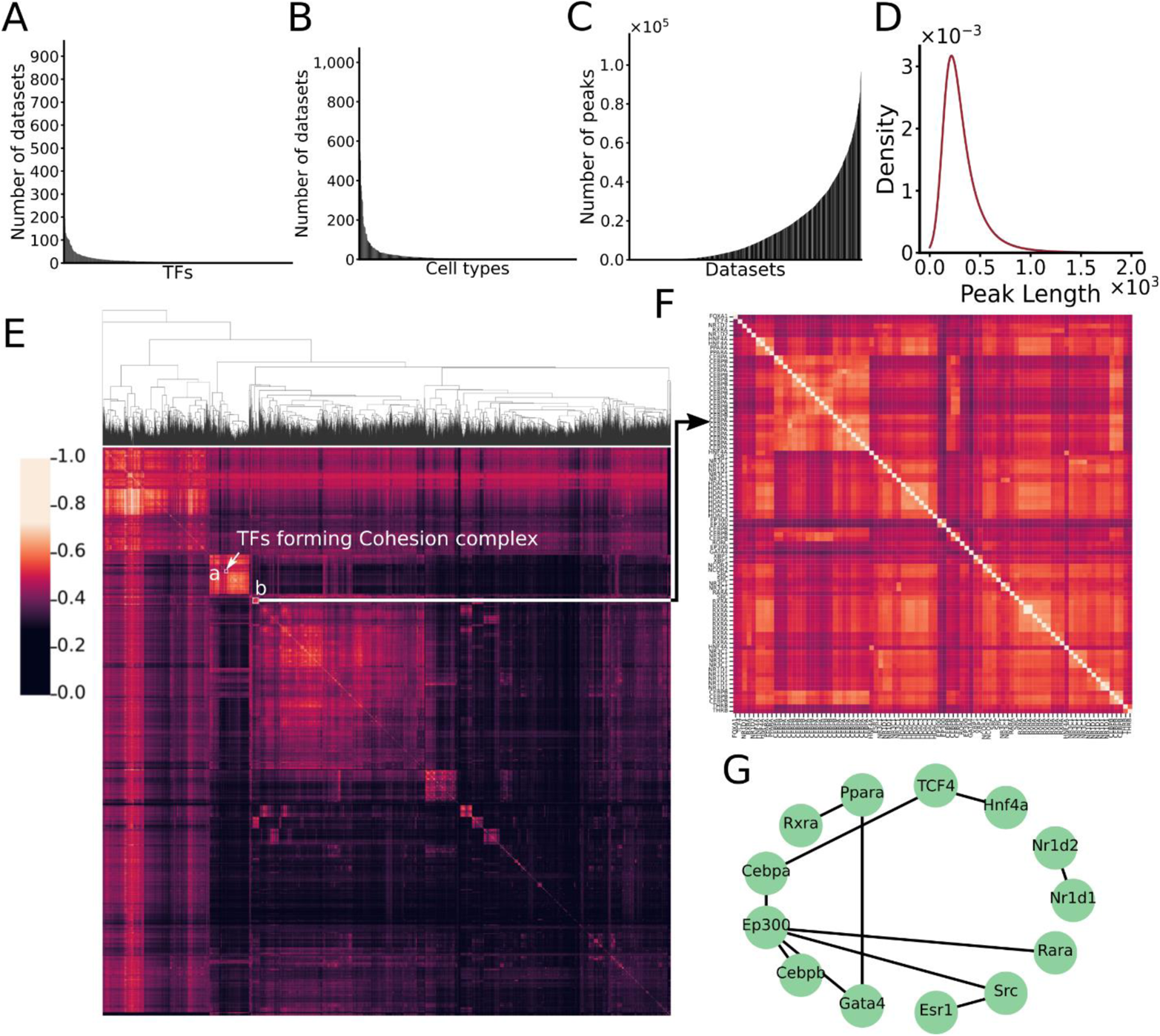
Evaluation of the TF ChIP-seq datasets. A. Number of datasets collected in each cell/tissue types sorted in the descending order. B. Number of datasets collected for each ChIP-ed TF sorted in the descending order. C. Number of peaks in each dataset sorted in the ascending order. D. Distribution of the lengths of originally called binding peaks in the 9,055 datasets. E. Heatmap of overlaps of the 1,000bp binding peaks between each pair of the datasets. The highlighted cluster *a* is formed by 50 datasets for Ctcf and varying number of datasets for its cooperative TFs Rad21, Stag1, and Smc1A in different cell/tissue types. F. A blowup view of cluster *b* highlighted in E, formed by 88 datasets for 20 TFs. G. Known physical interactions between 13 of the 20 TFs for which the 88 datasets form the cluster in F.

The DePCRM2 algorithm predicts CRMs and constituent TFBSs by identifying repeatedly cooccurring TFBSs of cooperative TFs in the 1,000bp binding peaks in all the collected ChIP-seq datasets for various TFs in different cell/tissue types of the organism (Niu et al. 2018; Ni and Su 2021). This design of the dePCRM2 algorithm was based on the observation that most cooperative TFs are often reused in various cell/tissue types at different developmental stages and/or under different homeostasis conditions (Davidson 2006). In theory, the larger the number of TF ChIP-seq datasets available and used, and the less bias of the datasets to few TFs and cell/tissue types, the better predictions that dePCRM2 can achieve (Niu et al. 2018; Ni and Su 2021). To see whether such highly biased datasets include enough datasets for cooperative TFs that are reused in different cell/tissue types, we calculated an overlapping score *S_o_* (formula 1) for each pair of the 9,055 non-empty datasets, and hierarchically clustered them. As show in Figure 1E, there are numerous overlapping clusters among the datasets which are either for largely the same TFs that were ChIP-ed in different cell/tissue types, or for different known cooperative TFs that were ChIP-ed in the same and/or different cell/tissue types. Interestingly, as seen in the human datasets (Ni and Su 2021), a cluster is formed by 50 datasets for cooperative TFs Ctcf, Rad21, Stag1, and Smc1A in various cell/tissue, reflecting the conserved cooperative relationships of the TFs in forming the cohesin complex (Zuin et al. 2014). Shown in Figure 1F is another example of cluster formed by 88 datasets for 20 TFs in various cell/tissue types, many of these TFs are known or likely to collaborate with each other according their physical interactions documented in the BioGRID (Oughtred et al. 2021) and reactome (Jassal et al. 2020) databases (Figure 1G). Therefore, notwithstanding these datasets are highly biased to few TFs (Figure 1A) and cell/tissue types (Figure 1B), they include datasets of many cooperative TFs that are reused in various cell/tissue types. The 1,000bp peaks in all the 9,055 datasets contain a total of 136,441,496,000bp, which is 50.1 times the size of the mouse genome (version mm10/GRCm38), but cover only 2,178,603,271bp (79.9%) of the mappable genome (2,725,521,370bp). Compared with the originally called peaks that cover a total of 1,398,035,305bp (51.3%) of the mappable genome, we substantially increased the coverage of the genome by extending the called peaks to 1,000bp, the size of shorter enhancers (Visel et al. 2007). dePCRM2 will predict which DNA segments in the 79.9% genome regions covered by the 1,000bp peaks are CRM candidate (CRMCs), and which are non-CRMCs, based on cooccurring patterns of putative TFBSs of motifs found in the binding peaks in all the datasets.

### Most of identified unique motifs (UMs) resemble known motifs and show intensive cooccurring pattens

dePCRM2 (Ni and Su 2021) starts by identifying all possible motifs in each dataset using ProSampler, an ultrafast motif finder (Li et al. 2019). ProSampler finds at least one motif in 8,294 (91.6%) of the 9,055 datasets, with a total of 1,062,339 motifs found. As shown in Figure 2A, the number of motifs found in a dataset increases with the number of peaks in it, but becomes stabilized around 250 when the number of peaks is above 50,000. dePCRM2 next identifies co-occurring motifs pair (CPs) as potential motifs, thereby filtering out most spurious motifs. To do so, dePCRM2 computes a co-occurring score *S_c_* (formula 2) for each pair of motifs in each dataset and selects the pairs with high scores as CPs. As in the case of human genome (Ni and Su 2021), the *S_c_* scores show a trimodal distribution (Figure 2B). dePCRM2 selects motifs pairs as PCs that account for the mode with the highest *S_c_* scores (Sc> 0.7 by default). More specifically, dePCRM2 identifies 4,028,221 CPs containing 225,809 (21.3%) potential motifs from 7,076 (85.3%) of the 8,294 datasets, while filtering out the remaining 1,218 (15.7%) datasets where no CPs are kept, and 836,530 (78.7%) possible spurious motifs. Many motifs in different CPs can be sub-motifs of the same TF, or of different members of a TF family that recognize highly similar motifs (Lambert et al. 2018; Ambrosini et al. 2020). Therefore, dePCRM2 clusters the 225,809 motifs in the 4,028,221 CPs by constructing a graph whose nodes are the motifs and edges are the SPIC similarity score (Zhang et al. 2013) between the motifs pairs, and then cutting the graph into dense subgraph as clusters of similar motifs. This results in 276 clusters, each containing from 28 to 49,308 motifs (Figure S1A). From these 276 motif clusters, dePCRM2 identifies 238 unique motifs (UMs) (Figure S1B). The UMs contain highly varying number of TFBSs, ranging from 72 to 14,025,382 with an average of 1,107,677 (Figure 2C). The lengths of the UMs range from 10 bp to 20 bp with a mean of 10.3bp, and are in the range of the lengths of known TF binding motifs (Figure 2D). The bias of the lengths of UMs to 10 bp is due to the limitation of ProSampler that needs to be improved. As expected, the UMs and their member motifs are highly similar to one another. For example, UM41 resembles its 11,799 highly similar member motifs (Figure 2E). To evaluate the UMs, we compared the 238 UMs against 875 annotated non-redundant motifs in the HOCOMOCO (Kulakovskiy and Makeev 2013; Kulakovskiy et al. 2018) and JASPER (Mathelier et al. 2016) databases using TOMOM (Gupta et al. 2007). Of the 238 UMs, 146 (61.3%) match at least one annotated motif, and 113 (77.4%) of the 146 UMs match at least two (Table S2), suggesting that most of the UMs might represent the motifs of the same TF family/superfamily which bind highly similar motifs (Lambert et al. 2018; Ambrosini et al. 2020). For instance, UM41 matches known motifs of five TFs of the “Jun-related factors” family (Jund, Bach1, Bach2, Junb and Nfe2) (Figure 2F), and five TFs of the “Fos-related factors” family (Atf3, Fosl2, Fosb, Fosl1 and Fos) (Table S2). On the other hand, the remaining 92 UMs might be novel motifs of unknown cognate TFs. We also evaluated the coverage of the UMs on motif families in the two databases (Mathelier et al. 2016; Kulakovskiy et al. 2018), and found that 82 (64.1%) of the 128 annotated TF motif families match one of the 238 UMs (Table S2), indicating that our predicted UMs recovery most of the known TF motif families.

**Figure 2.**
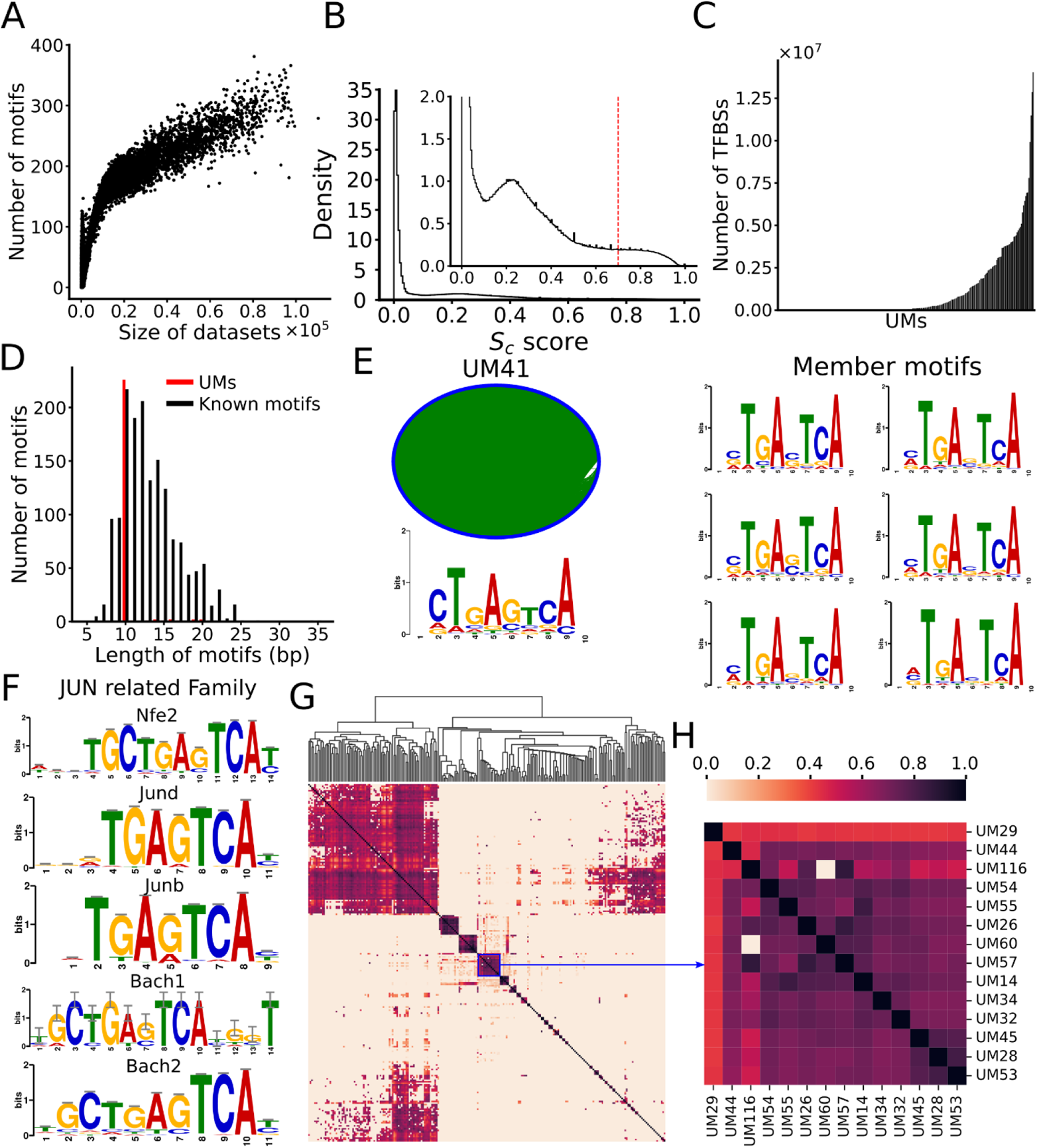
Prediction of UMs. A. Relationship between the number of predicted motifs in a dataset and the size (the number of binding peaks in the dataset). The datasets are sorted in the ascending order of their sizes. B. Distribution of cooccurrence scores (*S_c_*) of motif pairs found in each dataset. The dotted vertical line indicates the cutoff value of *S_c_* for predicting cooccurring pairs (CPs). C. Number of putative binding sites in each of the UMs sorted in the ascending order. D. Distribution of the lengths of the UMs and known motifs in the HOCOMOCO and JASPAR databases. E. The logo of UM41 and the motif similarity graph of its 11,799 member motifs that form a densely connected cluster. In the graph, each node in blue represents a member motif, and two member motifs are connected by an edge in green with SPIC score >0.8. Six examples of the member motifs are shown in the left panel. F. UM41 matches known motifs of five TFs of the JUN-related family. G. Heatmap of the cooccurrence/interaction networks of the 238 UMs, names of most UMs are omitted for clarity. H. A blowup view of the indicated cluster in G, formed by 14 UMs, of which UM116, UM14, UM26, UM28, UM29, UM32, UM45, UM53, UM55, and UM57 match known motifs (see main text).

To model cooccurring patterns of the UMs and interactions between their cognate TFs, dePCRM2 computes a cooccurrence/interaction score *S_INTER_* (formula 3) between each pair of UMs based on the co-occurrence of binding sites of UMs. As shown in Figure 2G, there are extensive cooccurrences between the UMs and interactions of their cognate TFs. These patterns of cooccurrences of the UMs indeed reflect the interactions among their cognate TFs or TF families for transcriptional regulation. For example, in a cluster formed by 14 UMs (Figure 2H), 10 of them (UM14, UM26, UM28, UM29, UM32, UM45, UM53, UM55, UM57 and UM116) match known motifs of TF families. More specifically, UM116 matches Msantd3, UM14 matches Ctcfl, UM26 matches Nfe2|Fosb|Atf3|Bach1|Pknox1|Jund|Nkx2-2|Jdp2|Fos|Junb|Fosl1|Fosl2|Batf|Msantd3|Bnc2|Mafk|Pbx3|Batf3|Jun, UM28 matches Zfp57|Atf3, UM29 matches Sp3|Mxi1|Nr1h4|Plagl1|Zfx|Klf3|Rfx1, and UM57 matches Nkx2-5|Fos|Fosb|Atf3|Pbx3|Junb|Jund|Pknox1|Fosl1|Batf3|Fosl2|Jun|Batf|Nkx2-2|Msantd3|Bnc2, etc. Some of these TFs are known collaborators in transcriptional regulation, such as Fos and Jun(Chevray and Nathans 1992; Norwitz et al. 2002; Miyamoto-Sato et al. 2005; de Marval et al. 2011), Atf3 and Jun(Yan et al. 2011), Pbx3 and Pknox1(Ravasi et al. 2010), Jun and Batf (Ravasi et al. 2010).

### Prediction of CRMs and constituent TFBSs in the mouse genome

To predict CRMs and constituent TFBSs in the mouse genome, dePCRM2 projects the TFBSs of the UMs to the genome and links adjacent TFBSs if their distance is less than 300bp (roughly, the length of two nucleosomes). dePCRM2 predicts each linked sequence as a CRM candidate (CRMC) and each sequence between two adjacent CRMCs in the peak-covered regions as a non-CRMC, thereby partitioning the peak-covered genome regions in two exclusive sets, CRMCs and non-CRMCs. Concretely, dePCTM2 predicts a total of 912,197 CRMCs and 1,270,937 non-CRMCs in the peak-covered genome regions, consisting of 55.5% and 24.4% of the genome, respectively. The CRMCs contains a total of 125,113,756 TFBSs, consisting of 23.9% of the genome and 42.9% of the CRMCs (Figure 3A). Many of these TFBSs have overlaps due partially to the aforementioned limitation of our motif-finder ProSampler, although it has been shown that certain patterns of transcriptional regulation are achieved by competitive or cooperative binding of the same or different TFs to overlapping TFBSs in a CRM (Xu et al. 1993; Drawid et al. 2009; Darieva et al. 2010; Inukai et al. 2017; Ostler et al. 2021). We connected each two adjacent overlapping putative TFBSs, resulting in a total of 38,554,729 non-overlapping putative TFBS islands with a mean length of 17bp.

**Figure 3.**
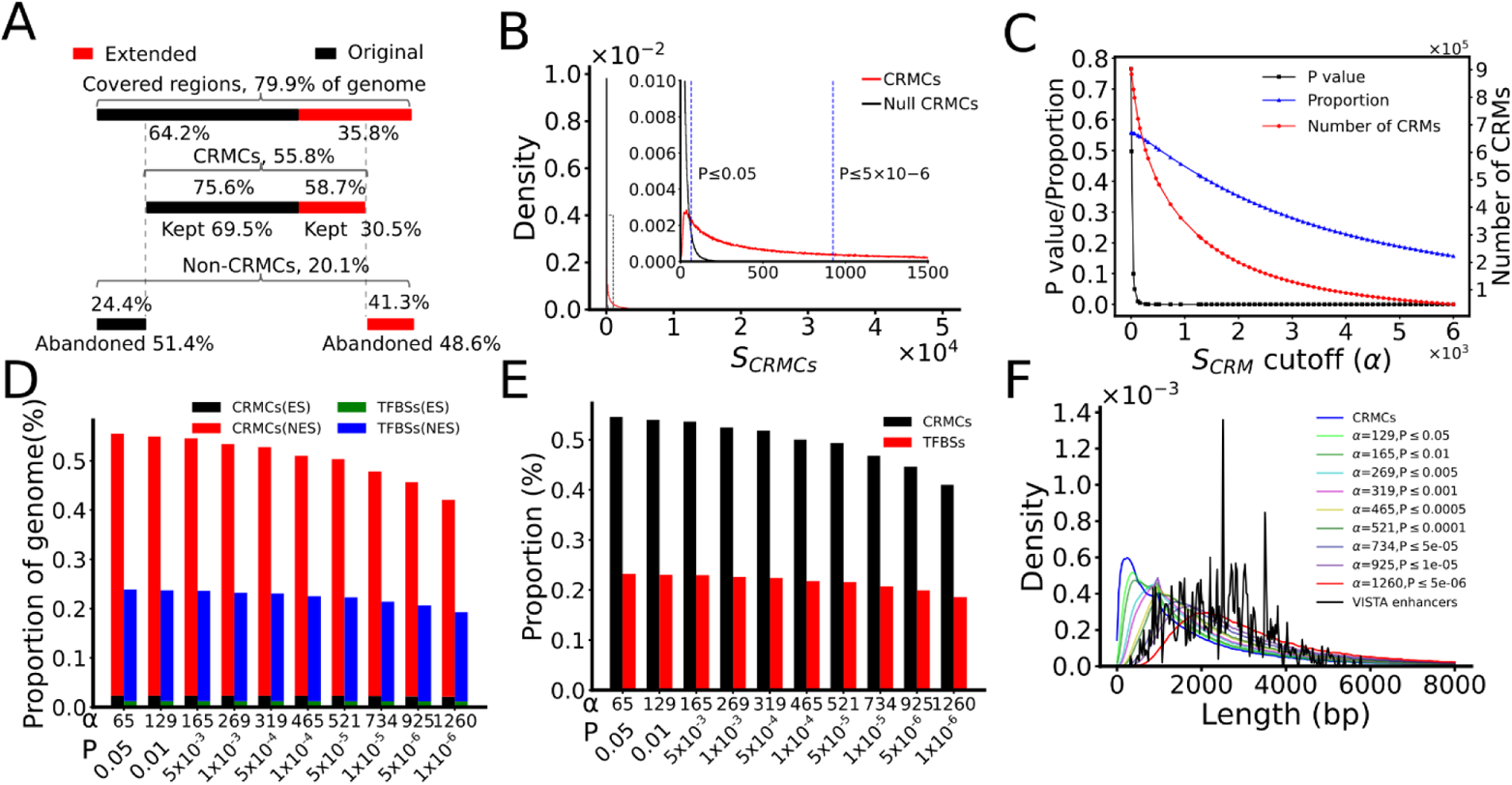
Prediction of CRMs using different *S_CRM_* cutoffs. A. A cartoon shows the proportions of the 79.9% of genome regions covered by originally called binding peaks (64.2%) and their extended parts (35.8%) as well as their relative contributions to the predicted CRMs (kept original (69.5%) and kept extended (30.5%)) and non-CRMCs (abandoned original (51.4%) and abandoned extended (41.3%)). Percentage above the lines are the proportion of originally called binding peak and their extended parts that are predicted to be CRMCs and non-CRMCs. B. Distribution of the *S_CRM_* scores of the CRMCs and the Null CRMCs. The inset is a blowup view of the indicated regions. The dotted vertical lines indicate *S_CRM_* cutoffs for the corresponding p-values. C. Number of the predicted CRMs, proportion of the genome predicted to be CRMs and the corresponding p-value as functions of the *S_CRM_* cutoff α. D. Percentage of the genome that are predicted to be CRM and TFBS positions in exonic sequences (ESs) and non-exonic sequences (NESs) using various *S_CRM_* cutoffs and corresponding p-values. E. Percentage of NESs that are predicted to be CRMs and TFBSs using various *S_CRM_* cutoffs and corresponding p-values. F. Distribution of the lengths of CRMs predicted using different *S_CRM_* cutoffs and corresponding p-values.

Interestingly, as in the case of human genome (Ni and Su 2021), 75.6% of genome positions of the originally called binding peaks were predicted as CRMC positions (kept-original), while the remaining 24.4% were predicted as non-CRMC position (abandoned-original) (Figure 3A). On the other hand, 58.7% of the extended positions were predicted as CRMCs (kept-extended), while the remaining 41.3% were predicted as non-CRMC positions (abandoned-extended) (Figure 3A). These results suggest that originally called binding peak positions may not necessarily parts of CRMs, while many flanking positions of the called peaks may be parts of CRMs. Therefore, as we concluded earlier (Ni and Su 2021), extension of the originally called peaks to roughly half of the mean length (1,000bp) of known of CRMs (2,400bp) (Visel et al. 2007) could greatly increase the chance of finding more CRMs in genomes.

To evaluate the CRMCs, dePCRM2 computes a *S_CRM_* score (formula 3) and a corresponding p-value for each CRMC (Materials and Methods). As shown in Figure 3B, the distribution of the *S_CRM_* scores of the CRMCs is strongly right-skewed relative to that of the Null CRMCs with the same number and lengths of the CRMs (Materials and Methods), suggesting that the CRMCs are unlikely produced by chance. Moreover, with the increase in the *S_CRM_* cutoff α, the corresponding p value drops rapidly, while both the number of predicted CRMs with a *S_CRM_* > *α* and their coverage of the genome decrease only slowly (Figure 3C), suggesting that most of the CRMCs have quite low p-values. More specifically, when the p-value drops precipitously from 0.05 to 1.00×10^−6^ the number of predicted CRMs and their coverage of the genome only decrease from 798,257 to 295,382, and from 55.5% to 42.1%, respectively (Figure 3D). Moreover, with the p-value dropping from 0.05 to 1.00×10^−6^, the coverage of putative TFBSs on the genome decreases only from 23.9% to 19.3%, and their percentage in the CRMs increases only from 43.0% to 45.8% (p-value ≤ 1.00×10^−6^) (Figure 3D). As expected, in the 0.05~1.00×10^−6^ range of p-value cutoffs, the vast majority of the predicted CRM positions (94.9~95.9%) and constituent TFBS positions (93.8~94.8%) are located in non-exonic sequences (Figure 3D), converging 41.0~54.70% and 18.6~23.2% of their lengths, respectively (Figure 3E). Interestingly, the remaining 4.1~5.1% of the predicted CRM positions and 5.2~6.2% of constituent TFBS positions are located in exonic sequences (Figure 3D), a well-known phenomenon in mammal genome (Mous et al. 1985; Yang et al. 1986; Farnham and Means 1990; Hurt et al. 1991; Hoeben et al. 1995; Neznanov et al. 1997; Chiquet et al. 1998; Lang et al. 2005; McLellan et al. 2006; Barthel and Liu 2008; Chen et al. 2008; Lampe et al. 2008; Tumpel et al. 2008; Dong et al. 2010; Birnbaum et al. 2012; Hirsch and Birnbaum 2015; Li et al. 2015). We will address these exonic CRMs and TFBS positions in great detail elsewhere.

We next compared the lengths of predicted CRMs at different *S_CRM_* cutoffs α with those of known mouse enhancers in the VISTA database (Visel et al. 2007). As shown in Figure 3F, the predicted CRMCs have a shorter mean length (1,682bp) than the VISTA enhancers (2,432bp). This is not surprising since most VISTA enhancers are involved in complex embryonic development and tend to be longer than other types of enhancers (Li and Wunderlich 2017). However, with the increase in the *S_CRM_* cutoff α, the distribution of the lengths of predicted CRMs shifts to right. Specifically, 252,349 (27.7%) of the 912,197 CRMCs were shorter than the shortest VISTA enhancer (330bp), but they cover only 2.1% of total length of the CRMCs, suggesting that they are likely either short CRMs or components of full-length enhancers remained to be fully predicted using more TF ChIP-seq datasets in the future. The remaining 659,848 (72.3%) CRMCs that are longer than the shortest VISTA mouse enhancer (330bp) consist of 97.9% of the total length of the CRMCs, and they are likely full-length CRMs. Thus, the vast majority (97.9%) of the CRMC positions are covered by predicted full-length CRMs. The predicted CRMs and constituent TFBSs are available at (https://cci-bioinfo.uncc.edu).

### Predicted CRMCs tend to be under strongly evolutionary constraints

To see how the CRMCs and non-CRMC evolve, we plotted the distributions of the phyloP scores (Pollard et al. 2010) of their nucleotide positions. The phyloP treats negative and positive selections in a unified manner and detects departures from the neutral rate of substitution in either direction, while allowing for clade-specific selection (Pollard et al. 2010). A positive phyloP score indicates the position is under purifying selection, a negative score indicates the position is under positive selection, and a score around zero means the position is selective neutral or nearly so. For convenience of discussion, we consider a position with a score in the range [−δ, δ] (δ>0) to be selectively neutral, in the range (δ, max) to be under positive selection, and in the range (min, −δ) to be under negative selection, respectively. We define the proportion of neutrality of a set of position as the areas under the distribution of the scores within range [−δ, δ], and choose δ= 1 in this study. For this analysis, we focused on the CRMCs and the non-CRMCs in non-exonic sequences, because including exonic sequences would confound the analysis due to their coding functions. The distribution of the phyloP scores of the non-CRMCs peaks at the neutral range with a proportion of neutrality of 0.89 (Figure 4A), suggesting that the non-CRMC positions are largely selective neutral as expected, although it is possible that some non-CRMC positions that are under some level of selections might have functions other than *cis*-regulatory. In contrast, the distribution of the phyloP scores of the CRMC positions displays a lower peak in the neutral range with a proportion of neutrality of 0.77 (Figure 4A), and spreads to both negative selection and positive selection ranges. These results indicate that CRMC positions are more likely to be under evolutional constraints than the non-CRMC positions. Thus, the CRMCs are more likely to be functional than non-CRMCs, although some CRMC positions that are selected neutral might not be functional. Notably, the mouse VISTA enhancers are even more likely to be evolutionarily conserved than our predicted CRMCs (Figure 4A), although the former are largely a small subset of the latter (see below). This is not surprising that the VISTA enhancers were selected for validation in transgene animal models due to their ultra-conservation (Visel et al. 2008) and thus are mainly involved in embryonic development (Bejerano et al. 2004; Katzman et al. 2007) Therefore, as in the case of the human genome (Ni and Su 2021), dePCRM2 is able to partition the peak-covered genome regions into a functional set, i.e., the CRMCs, and a non-functional set, i.e., the non-CRMCs.

**Figure 4.**
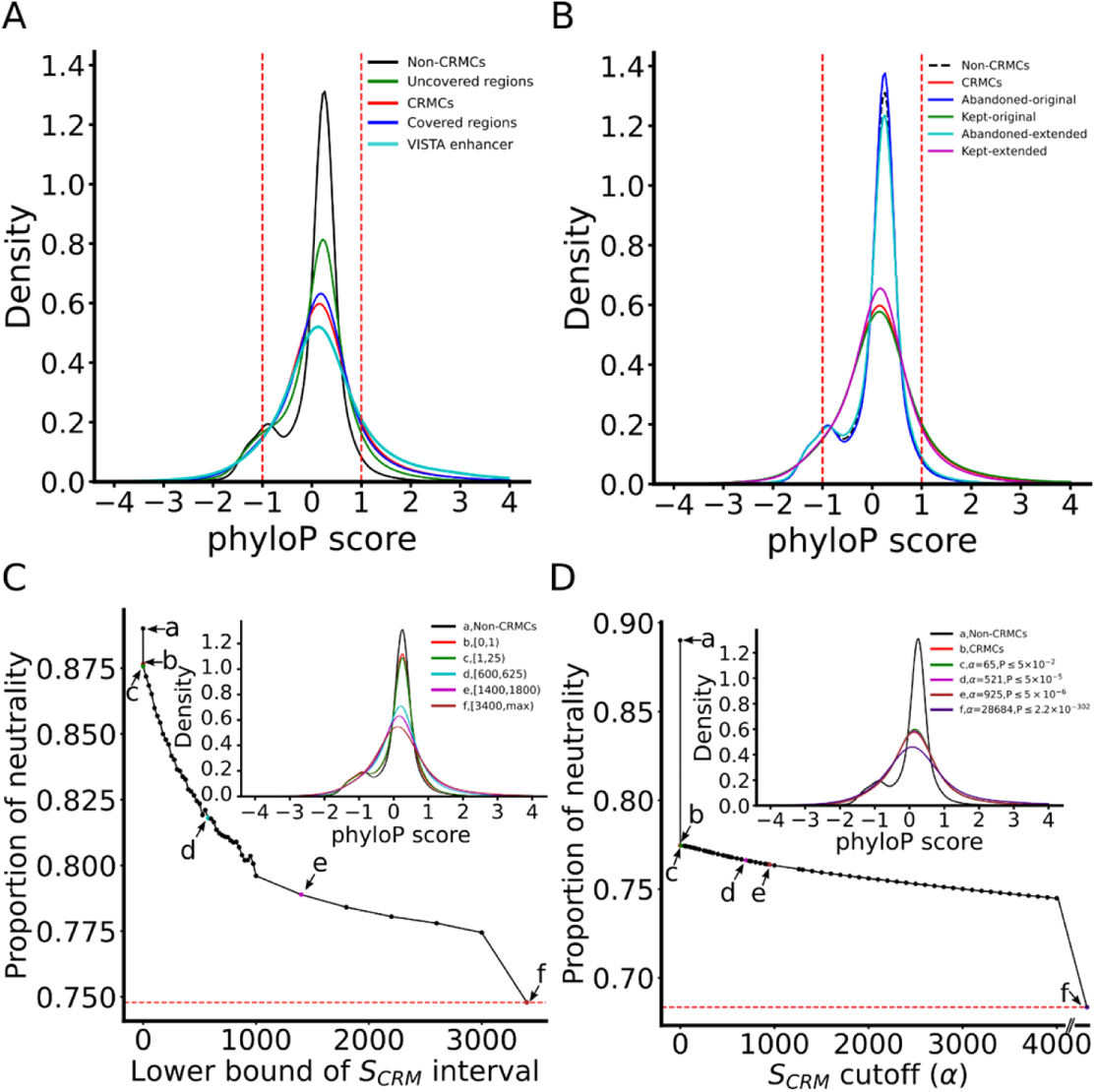
Different evolutionary constraints on the predicted CRMCs and the non-CRMCs in non-exonic sequences measured by phyloP scores. A. Distributions of phyloP scores of nucleotide positions of the VISTA enhancers, the predicted CRMCs, the non-CRMCs, peak-covered regions and peak-uncovered regions. The area under the density curves in the score interval [−1, 1] is defined as the proportion of neutrality of the positions. B. Distributions of phyloP scores the kept-original, the kept-extended, the abandoned-original and the abandoned-extended positions in comparison with those of the CRMCs and the non-CRMCs. The distributions for the kept-original positions and the kept-extended positions are significantly different from those of the abandoned-original positions and the abandoned-extended positions, respectively, p<2.2×10^−302^ (K-S test). C. Proportion of neutrality of the CRMCs with a *S_CRM_* score in different intervals in comparison with that of the non-CRMCs (a). The inset shows the distributions of the phyloP scores of the non-CRMCs and the CRMCs with *S_CRM_* scores in the intervals indicted by color and letters. D. Proportion of neutrality of the CRMs predicted using different *S_CRM_* score cutoffs and corresponding p-values in comparison with those of the non-CRMCs (a) and the CRMCs (b). The inset shows the distributions of the phyloP scores of the non-CRMCs, the CRMCs and the CRMs predicted using the *S_CRM_* score cutoffs and corresponding p-values indicated by color and letters.

As we indicated earlier, there are still 20.1% of genome regions that are not covered by the extended peaks. To see whether the non-exonic sequences in these peak-uncovered regions contain functional elements such as CRMs, we plotted the distribution of the phyloP scores of their genomic positions. The proportion of neutrality (0.83) of these positions is in between those of the peak-covered regions (0.78) and those of the non-CRMCs (0.89) (Figure 4A), suggesting that they might contain functional elements, albeit with a lower density than that in the peak-covered regions. Based on the difference in the proportion of neutralities of the peak-covered and peak-uncovered regions as well as that of the non-CRMCs, we estimate that proportion of CRMC positions in the peak-uncovered regions is about [(1-0.83)-(1-0.89)]/[(1-0.78)-(1-0.89)]=54.55% that of CRMC positions in the peak-covered regions.

As expected, the kept-original positions as well as the kept-extended positions have almost the same phyloP score distributions as the CRMCs (Figure 4B), indicating that they all are under strongly evolutionary constraints. In contrast, the abandoned-original peak positions as well as the abandoned-extended positions have an almost identical phyloP score distributions to that of the non-CRMCs (Figure 4B), indicating that they all are largely selectively natural or nearly so. These results strongly suggest that the kept extended positions are likely functional, while the abandoned-original positions are unlikely functional. This results confirm our earlier conclusion that originally called binding peaks cannot be equivalent to CRMs, and appropriate extension of the originally called short binding peaks can greatly increase the power of available datasets for predicting CRMs and constituent TFBSs in genomes (Ni and Su 2021).

### Higher-scoring CRMs are more likely under evolutionary constraints

To investigate the relationship between the evolutionary behaviors of the CRMCs and their *S_CRM_* scores, we plotted the distribution of the phyloP score of subsets of CRMCs with *S_CRM_* scores in nonoverlapping intervals. As shown in Figure 4C, with the increase in the *S_CRM_* scores, the proportion of neutrality of the corresponding CRMs first drops rapidly and then enters a gradually decreasing phase. Thus, CRMCs with higher *S_CRM_* scores are more likely under evolutionary constraints, indicating that the *S_CRM_* score captures the evolutionary behavior of a CRM. Interestingly, even the CRMCs with scores in the lowest interval [0, 1) have a lower proportion of neutrality than that of the non-CRMCs (0.87 vs 0.89) (Figure 4C), suggesting that even these lowest scoring CRMCs that tend to be short (Figure 3F) are under stronger evolution constraints than the non-CRMCs, and thus are likely functional.

Next, we examined the phyloP scores for the CRMs predicted at different *S_CRM_* score cutoffs α (or p-values). As shown in Figure 4D, with the increase in the *S_CRM_* score cutoff α, the proportion of neutrality of the predicted CRMs decreases gradually, suggesting again that the *S_CRM_* score captures the evolutionary behavior of the CRMCs. As indicated earlier, even at the lowest *S_CRM_* cutoff (α=0), the predicted CRMs (i.e., all the CRMCs) have smaller neutral composition than that of the non-CRMCs, suggesting that at least most of the CRMC are functional, and the higher the *S_CRM_* score of a CRM, the more likely it is evolutionarily constrained, and thus the more likely it is functional.

### Predicted CRMs are supported by independent experimental data

We next evaluated the sensitivity (recall rate) of our CRMs predicted at different p-values for recalling four types of experimentally determined CRM-related elements, including 620 mouse enhancers documented in the VISTA database (Visel et al. 2007), 163,311 mouse promoters and 49,385 mouse enhancers determined by the FANTOM project (Forrest et al. 2014; Lizio et al. 2019), and 2,208 QTLs documented in the Mouse Genome Informatics (MGI) databases (Bult et al. 2019). Interestingly, most of these experimentally determined elements are located in the peak-covered genome regions, including 579 (93.4%) VISTA enhancers, 163,311 (99.1%) FANTOM promoters and 49,385 FANTOM enhancers (99.2%) (Andersson et al. 2014), with the exception for QTLs with only 1,023 (46.3%) being located in the peak-covered regions. If a predicted CRM and an element overlaps each other by at least 50% of the length of the shorter one, we say that the CRM recovers the element. As show in Figure 5A, with the increase in the p value (decrease in −log(p)) cutoff, the sensitivity increases rapidly and saturates at a p-value cutoff 0.05 (α ≥ 65) to 99.3%, 93.8%, 86.8%, 82.3% for recovering the VISTA enhancers, FANTOM promoters, FNATOM enhancers and QTLs, respectively. Thus, the VISTA enhancers are largely a subset of our CRMCs. In contrast, the control sequences with the matched number and lengths of the predicted CRMs at different p-value cutoffs only recall an expected proportions of the elements by chance (p<2.2×10^−302^, χ^2^ test) (Figure 5A). Figures S2A~S2D show examples of the predicted CRMs that recover these four different types of experimentally determined elements.

**Figure 5.**
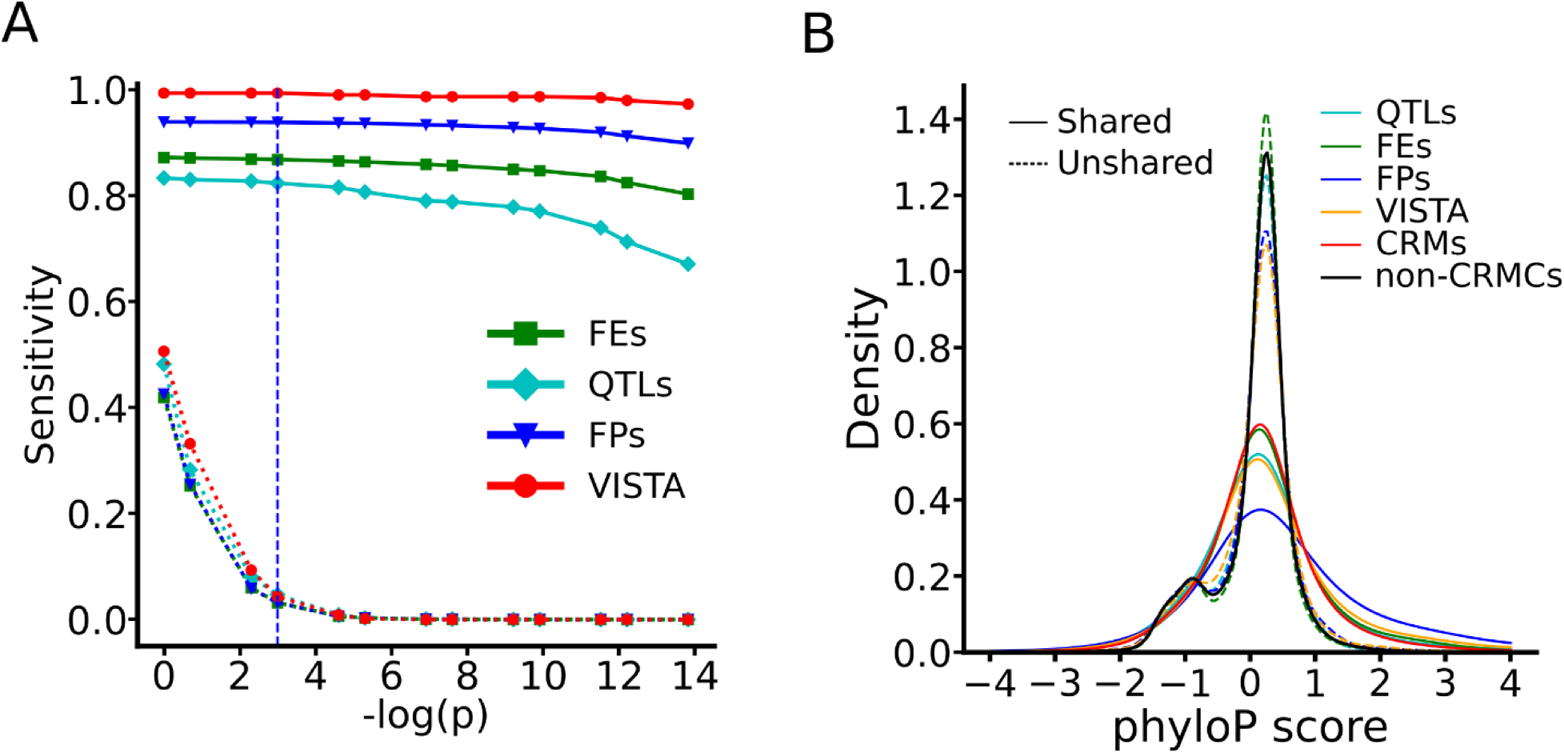
Validation of the predicted CRMs by VISTA enhancers, FANTOM promoters (FPs), FANTOM enhancers (FEs) and QTLs. A. Sensitivity (recall rate) of the predicted CRMs or the control sequences as a function of p-value cutoff for recalling each set of the experimentally determined elements. The dashed vertical line indicates the p-value cutoff of 0.05. The sensitivity of the CRMs predicted at all the indicated p-value cutoffs are significantly higher (p<2.2×10^−302^, χ^2^ test) than the control sequences for recalling each set of the experimentally determined elements. B. Distributions of phyloP scores of the shared and unshared nucleotide positions of the elements in each set of the experimentally determined elements, in comparison with those of the predicted CRMs at p≤0.05 and of the non-CRMCs. The difference between the distributions of shared and unshared positions in each set of the experimentally determined elements is significant, p<2.2×10^−302^ (K-S test). Note that there are only three unrecalled VISTA enhancers.

The varying range of sensitivity from 82.3% for QTLs to 99.3% for VISTA enhancers might reflect the varying reliability of methods used to characterize these four types of elements. For example, VISTA enhancers and FANTOM promoters were determined by highly reliable transgene animal models (Visel et al. 2007) and CAGE methods (Lizio et al. 2015), respectively, and our predicted CRMs achieve very high sensitivity to recall them. On the other hand, FANTOM enhancers and QTLs were determined by less reliable eRNA quantification (Andersson et al. 2014) and association studies, respectively, and our predicted CRMs achieve relatively low sensitivity to recall them.

To find out whether our predicted CRMs missed these unrecalled elements, or they are simply false positives due to the limitations of experimental methods used to characterize them, we compared the phyloP scores of the recalled and unrecalled elements. As shown in Figure 5B, for all the four types of elements, the recalled elements (solid lines) tend to be under strongly evolutionary constrains like our predicted CRMs, thus are likely functional. In contrast, the unrecalled elements (dashed lines) are largely selective neutral like our predicted non-CRMCs, thus are likely false positives produced by the methods used to characterize them. Based these results, we estimated an FDR of 0.7% (100%-99.3%), 6.2% (100%-93.8%), 13.2% (100%-86.8%) and 17.7% (100%-82.3%) in VISTA enhancers, FANTOM promoters, FANTOM enhancers and QTLs, respectively.

### Most of predicted CRMs might be in correct lengths

Correct characterization of the lengths of CRMs is notoriously difficult both experimentally and computationally, because even short components of a long CRM might still be at least partially functional in transgene animal models (Visel et al. 2008), and because functionally related independent enhancers may cluster with each other to form super-enhancers (Pott and Lieb 2014; Dukler et al. 2016), or locus control regions (LCRs) (Li et al. 2002). Although VISTA enhancers are by no means a gold standard set of CRMs with correctly characterized lengths (Visel et al. 2007), they are the only available set of validated enhancers in mouse. As we indicated earlier, our CRMs predicted at p-value cutoff 0.05 recall 575 (99.3%) of the 579 VISTA enhancers in the peak-covered genome regions (Figure 5A), we thus ask whether the recalling CRMs have a length matching the recalled VISTA enhancers. To this end, we computed the ratio of the length of a recalling CRM over that of its recalled VISTA enhancer. As shown in Figure 6A, the recalling CRMs are on average twice as long as the recalled VISTA enhancers. To see whether we over-predict the lengths of the recalling CRMs or the recalled VISTA enhancers are only shorter functional components of long enhancers, we compared phyloP scores of the 1,303,562bp positions shared by the recalling CRMs and the recalled VISTA enhancers, with those of the 3,005,862bp (69.75%) and 73173bp (5.31%) positions specific to the recalling CRMs and the recalled VISTA enhancers (Figure 6B). As expected, like our predicted CRMC positions (Figure 4A), positions shared by the CRMs and the VISTA enhancers tend to be under strongly evolutionary constraints (Figure 6C). Moreover, the CRM specific positions (69.75%) also tend to be under strongly evolutionary constraints (Figure 6C) as expected, suggesting that the positions in recalling CRMs that the recalled VISTA enhancers lack might be functional. In contrast, like our predicted non-CRMCs (Figure 4A), the VISTA enhancer specific positions (5.31%) are largely selectively neutral (Figure 6C), suggesting that the positions in the recalled VISTA enhancers that the recalling CRMs lack might not be functional. Therefore, although the recalled VISTA enhancers are only half as long as the recalling CRMs, they might contain non-enhancer sequences that comprise 5.31% of the total length of the recalled VISTA enhancers. On the other hand, we noted that 38 (6.6%) VISTA enhancers were recalled by multiple short CRMCs, suggesting that some of short CRMCs are indeed only components of a long CRMC, whose full-length forms remain to be predicted when more TF ChIP-seq data are available in the future.

**Figure 6.**
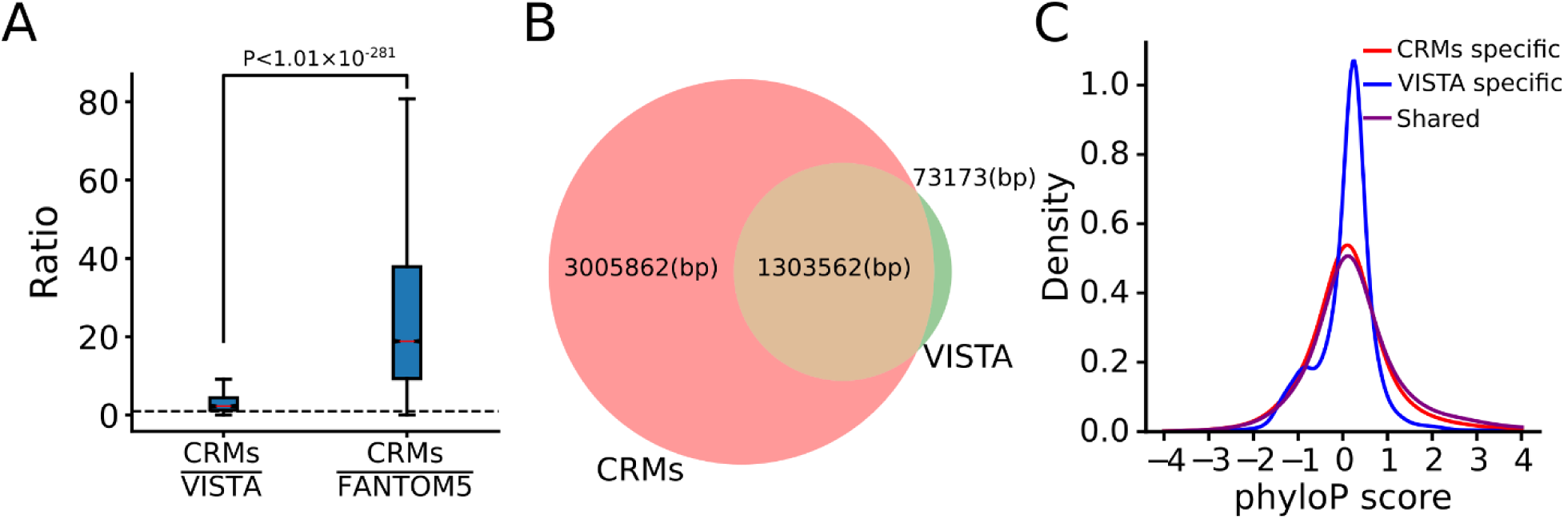
dePCRM2 might correctly predict the lengths of most CRMs. A. Boxplots of the ratio of the length of a recalling CRM over that of its recalled VISTA enhancers and FANTOM5 enhancers. The p value was calculated using the Mann-Whitney U test. B. Venn diagram showing the number of nucleotide positions shared by recalling CRMs and recalled VISTA enhancers, and the number of positions specific to the recalling CRMs and the recalled VISTA enhancers. C. Distributions of phyloP scores of the positions shared by the recalling CRMs and the recalled VISTA enhancers, and of the positions specific to the recalling CRMs and to the recalled VISTA enhancers. The difference between the distributions of the shared and VISTA specific positions is significantly different, p<2.2×10^−302^, K-S test.

We also compared the lengths of the recalling CRMs and their recalled FANTOM5 enhancers. As shown in Figure 6A, the recalling CRMs (median length 4,233bp) are about 14.7 times as long as the recalled FANTOM enhancers (median length 288bp). Moreover, 34.8% of the recalled FANTOM enhancers were located in the same CRMs. Thus, FANTOM enhancers tend to be short components of long CRMs. Taken together, these results strongly suggest that although some of our CRMCs might be short components of long CRMs, the vast majority of the CRMs predicted p-value cutoff of 0.05 might be in correct full length, while many VISTA enhancers and most FANTOM enhancers might be only a component of otherwise long enhancers.

### Our predicted CRMs and constituent TFBSs are more accurate and complete than existing predictions

We further evaluated our 798,257 CRMs predicted at p-value ≤ 0.05 (*S_CRM_* ≥ 65) with two sets of predicted mouse enhancers, including 339,815 cCREs predicted recently by the ENCODE phase 3 consortium (Moore et al. 2020) and 519,386 enhancers from the EnhancerAtlas database (Gao and Qian 2020). As shown in Figure 7A, these three sets of predicted CRMs containing highly varying numbers of elements cover highly varying portions of the genomes, i.e., 55.5%, 3.4% and 81.6% by our CRMs, the cCREs and the EnhancerAtlas enhancers, respectively. Since all our CRMs are located in the peak-covered genome regions, we only consider for comparison the cCREs and the EnhancerAtlas enhancers that have at least one nucleotide position overlapping the peak-covered genome regions. As shown in Figure 7A, the vast majority of the cCREs (339,721 or 99.97%) and the EnhancerAtlas enhancers (436,504 or 84.0%) have at least one nucleotide position overlapping the peak-covered genome regions. The cCREs and EnhancerAtlas enhancers that at least partially overlap the peak-covered genome regions cover 3.4% and 81.6% of the genome (Figure 7A). Therefore, our CRMs in the peak-covered genome regions cover a much larger proportion (55.5%) of the genome than do the cCREs (3.4%), but a much smaller proportion of the genome than do the EnhancerAtlas enhancers (81.6%).

**Figure 7.**
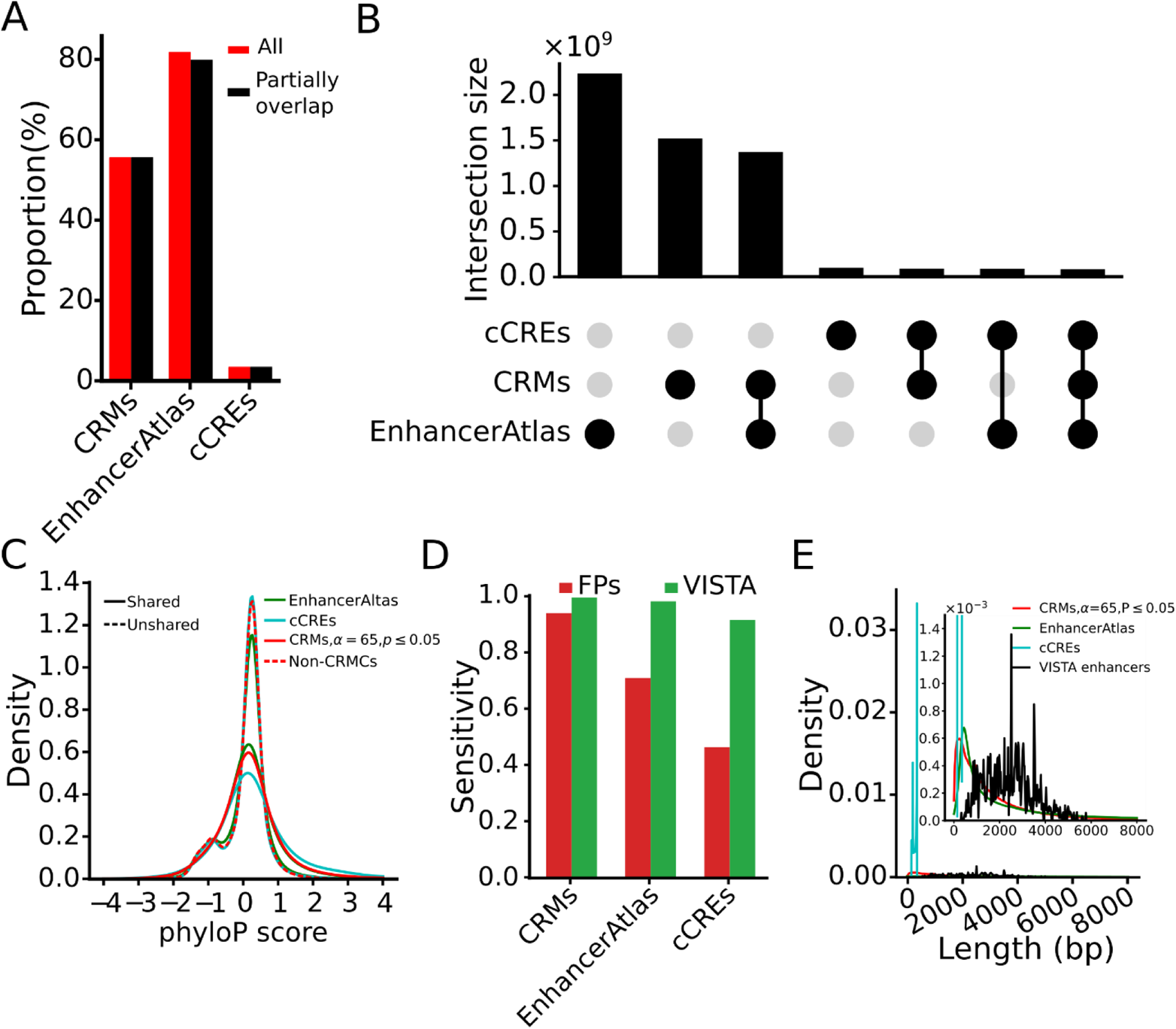
Comparison of our CRMs (p-value <0.05) with the cCREs and the EnhancerAtlas enhancers. A. Percentage of the genome covered by all the sequences of the CRMs, the cCREs and the EnhancerAtlas enhancers (All), and by the sequences in the CRMs, the cCREs and the EnhancerAtlas enhancers, which at least partially overlap the peak-covered genome regions (Partially overlap). B. Upset plot showing numbers of nucleotide positions shared and unshared among the sequences in the three sets of predicted CRMs. C. Distributions of phyloP scores of nucleotide positions of the cCREs and the EnhancerAtlas enhancers that are shared and unshared with our CRMs (p-value ≤ 0.05). D. Comparison of sensitivity of the three sets of predicted CRMs for recalling FANTOM promoters and Vista enhancers. E. Distributions of lengths of the CRMs, EnhancerAtlas enhancers and cCREs. The inset is a zooming-in view of the indicated range of the vertical axis.

To see whether we over-predicted the CRMs with respect to the cCREs, or under-predicted the CRMs with respect to the EnhancerAtlas, we first identified the shared and unshared genome positions among the three sets of sequence elements. As shown in Figure 7B, most (85,075,038bp or 92.0%) of the cCRE positions overlap our CRM positions, but they only cover 5.6% of our CRM positions, while missing 94.4% of our CRM positions because of the much shorter total lengths of the cCREs (Figure 7A). The remaining 8.0% of the cCRE positions do not overlap our CRM positions. A total of 1,364,995,621bp (61.3%) EnhancerAtlas enhancer positions overlap our CRM positions (Figure 7B), covering 90.3% of our CRM positions, while missing 9.7% of our CRM positions. The remaining 39.7% of the EnhancerAtlas enhancer positions do not overlap our CRMs.

We then compared the phyloP scores of the cCRE and EnhancerAtlas enhancer positions that they shared and unshared with our CRMs positions (Figure 7B). As expected, like our CRM positions (Figure 7C), both the cCRE and the EnhancerAtlas positions shared with our CRMs tend to be under strongly evolutionary constraints, suggesting that they are likely functional. In stark contrast, the eCRE and the EnhancerAtlas positions unshared with our CRMs are largely selectively neutral like the non-CRMCs, suggesting that they might be not functional, and thus are false positive predictions. These results suggest that the cCRE and EnhancerAtlas enhancer positions that overlap our CRMs are more likely to be functional, while those that do not overlap our CRMs are more likely to be false positives. Therefore, based on the proportion of the unshared positions, we estimate the FDRs of the cCREs and EnhancerAtlas enhancers to be about 8.0% and 39.7%, respectively.

We also compared sensitivity of our CRMs, EnhancerAtlas and cCREs for recalling FANTOM prompters and VISTA enhancers in the peak-covered genome regions. We choose the FANTOM promoters and VISTA enhancers for this validation because the high quality of the two datasets with an estimated FDR of 0.7% and 6.2%, respectively, based on their proportions of neutrality (Figure 5B). As shown in Figure 7D, our CRMs substantially outperform the cCREs for recalling the FANTOM promoters (93.8% vs 46.3%) and VISTA enhancers (99.3% vs 91.4%). However, this comparison might not be meaningful as the total length of our CRMs is 16 time as large as that of the cCREs. On the other hand, although the total length of our CRMs is only 68.0% that of the EnhancerAtlas enhancers, our CRMs outperform the EnhancerAtlas enhancers for recalling VISTA enhancers (99.3% vs 97.9%) and FANTOM promoters (93.8% vs 70.1%).

Finally, we compared the lengths of our CRMs with those of the cCREs and the EnhancerAtlas enhancers. As shown in Figure 7E, the distribution of the lengths of the cCREs has a very sharp peak around 250bp with a mean length of 272bp, indicating that the cCREs have almost the same lengths, a possible artifact of the prediction methods. Both the distributions of the lengths of our CRMs and EnhancerAtlas enhancers are similarly strongly skewed toward right with a mean length of 1,893 and 4,285bp, respectively. Since there is no gold standard set of full-length CRMs, we could not validate the length of our CRMs and EnhancerAtlas enhancers. However, based on the evolutionary constraints on our CRMs, most our predicted CRMs might be in full-length, while 39.7% of the EnhancerAtlas enhancers positions might be false positives as we argued earlier. Taken together, our results suggest that our CRMs might be more accurate and complete than both the cCREs and the EnhancerAtlas enhancers.

### About 64% of the mouse genome might code for CRMs

As we indicated earlier, our predicted 912,197 CRMCs make up of 55.5% of the mappable mouse genome. To estimate the FDR of the CRMCs, we took a semi-theoretic approach as we did earlier in the human genome (Ni and Su 2021). Specifically, we calculated the expected number of true positives and false positives in the CRMCs with a *S_CRM_* score in each of non-overlapping interval based on the density of the *S_CRM_* scores of the CRMCs and the density of the *S_CRM_* scores of the Null CRMCs (Figure 8A), yielding 910,711 (99.84%) expected true positives and 1,486 (0.16%) expected false positives in the CRMCs (Figure 8B). Most (1,373/1,486=92.40%) of the 1,486 expected false positive CRMCs have a low *S_CRM_* score < 50 (insets in Figures 8A and 8B) with a mean length of 64bp, comprising 0.004% (1,486*64bp/2,725,521,370bp) of the mappable genome and 0.007% (0.004/55.5) of the total length of the CRMCs, i.e., an FDR of 0.007% for the CRMC positions (Figure 8C). Thus, our predicted true CRMCs would comprise 55.5%-0.004%=55.496% of the genome. On the other hand, as the CRMCs miss 0.7% of VISTA enhancers in the peak-covered regions [the point at −log (p) =0 in Figure 5A], we assume the FNR of predicting CRMC positions to be about 0.7%. We estimate false negative CRMC positions to be 0.007*0.55496/(1-0.007)=0.39% of the genome, which is 0.39%/24.4%=1.60% of the total length of the non-CRMCs, meaning a false omission rate (FOR) of 1.60% for the non-CRMC positions (Figure 8C). Hence, true CRM positions in the peak-covered regions would be 55.5%-0.004%+0.39%=55.89% of the genome (Figure 8C). In addition, as we argued earlier, the CRMC density in the peak-uncovered 20.10% genome regions is about 54.55% of that in the peak-covered genome regions, CRMCs in the uncovered regions would be about 0.201*0.5589*0.5455/0.779=7.87% of the genome (Figure 8C). Taken together, we estimated about 55.89%+7.87%=63.76% of the genome to code for CRMs, for which we have predicted 55.89/63.76=87.66%. Moreover, as we predicted about 42.9% of CRMs to be made up of TFBSs (Figure 3D), we estimated about 0.429*63.76%=27.35% of the genome to encode TFBSs. Furthermore, assuming a mean length 1,893bp for CRMs (the mean length of our predicted CRMs at p-value ≤0.05), and a mean length of 17 bp for TFBS islands, we estimated that the mouse genome would encode about 918,010 CRMs (2,725,521,370x0.6376/1,893) and 43,848,829 non-overlapping TFBS islands (2,725,521,370x0.2735/17).

**Figure 8.**
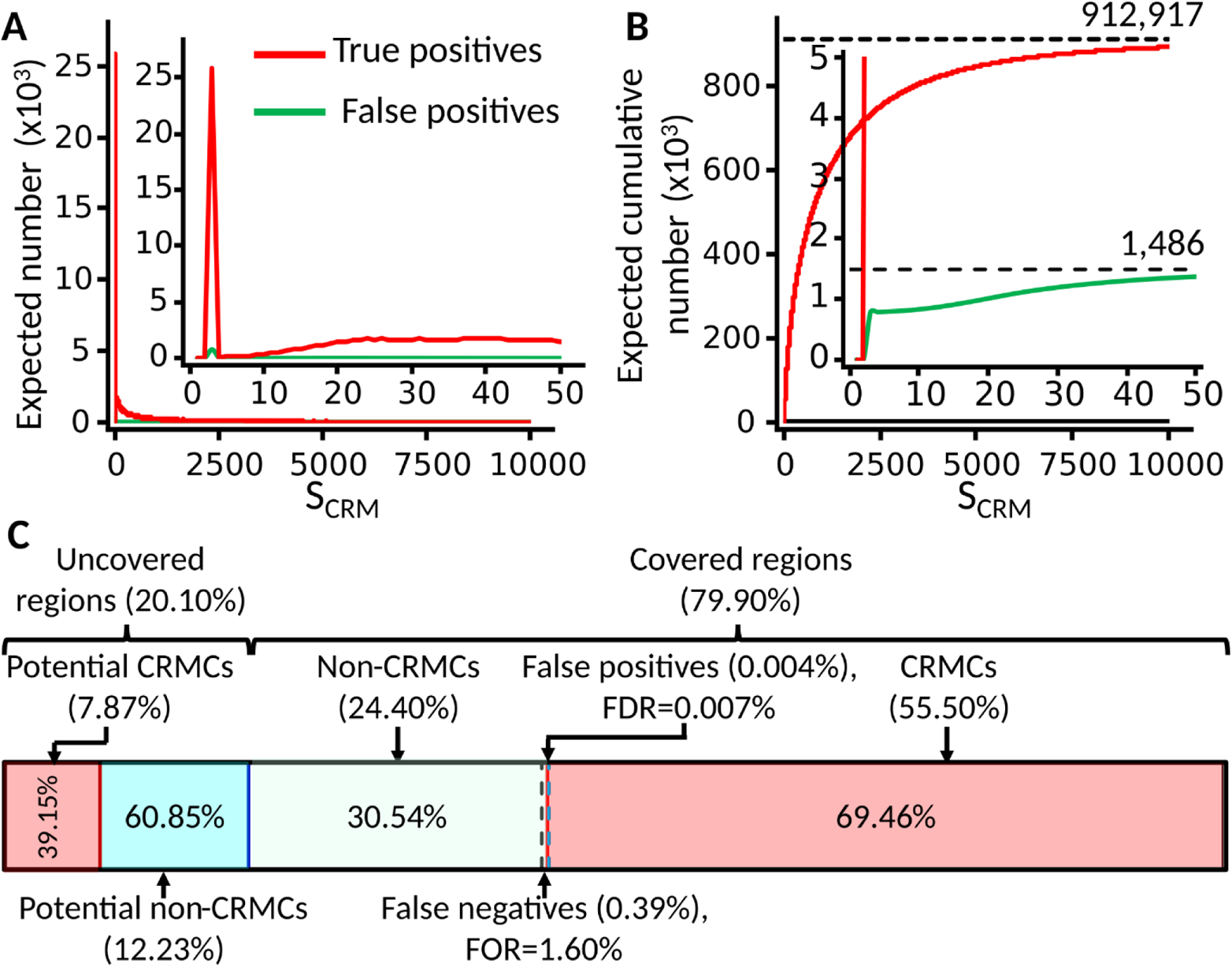
Estimation of the portion of the mouse genome encoding CRMs. **A.** Expected number of true positive and false positive CRMCs in the predicted CRMCs in each one-unit interval of the *S_CRM_* scores. The inset is a blow-up view of the axes defined region. **B.** Expected cumulative number of true positives and false positives with the increase in *S_CRM_* score cutoff for predicting CRMs. The inset is a blow-up view of the axes defined region. **C.** Proportions of the peak-covered genome regions (79.9%) and peak-uncovered genome regions (20.1%) in the genome and estimated proportions of CRMCs in them. Percentages in the braces are the proportions of the indicated sequence types in the genome, and percentages in the boxes are the proportions of the indicated sequence types in the covered regions or in the uncovered regions.

## DISCUSSION

In this study, using the dePCRM2 pipeline (Ni and Su 2021), we predicted an unprecedented comprehensives map of 0.91M CRMCs and 38.55M constituent TFBS islands in 79.9% of the mouse mappable genome covered by 1,000bp binding peaks in 9,060 ChIP-seq datasets for 701 TFs in 438 mouse cell line/tissue/organ types. Many features of the predicted CRMCs and TFBSs in the mouse genome are reminiscent of those of our earlier predicted CRMCs and TFBSs in the human genome (Ni and Su 2021). First, the number of predicted UMs in both genomes are very close (238 vs 210), reflecting the fact that both genomes encode highly conserved sets of TF families (Fulton et al. 2009; Lambert et al. 2018). Second, most of the UMs in both genomes match known TF motif families, and most known motif families are matched by the UMs in both genomes. Third, the mouse CRMCs consist of 55.5% of the mouse genome, while the human CRMCs make up of 44.0% of the human genome (Ni and Su 2021). The higher genome coverage of the mouse CRMCs are clearly due to a larger number (9,060 vs 6,092) of TF ChIP-seq datasets covering a higher proportion (79.9% vs 77.5%) of the mouse genome were used. Fourth, peak-uncovered regions in both genome may still contain CRMs albeit at a lower density than the peak-covered regions according to their evolutionary profiles (Figure 4A) (Ni and Su 2021). To predict CRMs and constituent TFBSs in these peak-uncovered regions in both genomes, more TF ChIP-seq data, particularly, for new TFs in new cell/tissues of human and mouse are needed to cover these currently peak-uncovered regions. We expect that with more TF ChIP-seq datasets available in both the human and mouse cell/tissue types, the peak-covered genome regions would increase and eventually become saturated (Niu et al. 2018; Ni and Su 2021). Fifth, we estimated that about 63.8% (Figure 8C) and 55.4% (Ni and Su 2021) of the mouse and human genomes might encode CRMs, and TFBSs make up of about 40% of the lengths of the CRMs in both genomes. Therefore, CRMs might be more prevalent than originally thought in both the mouse and human genomes. However, they might not be as prevalent (81% and 59% in the mouse and human genomes, respectively) (Figure 7C) (Ni and Su 2021) as the EnhancerAtlas database documented (Gao and Qian 2020).

Sixth, the predicted CRMCs in both genomes are more likely subject to evolutionary constraints than the predicted non-CRMCs that are largely selectively neutral or nearly so. Hence, the CRMCs are likely *cis*-regulatory, while the non-CRMCs are unlikely *cis*-regulatory. Seventh, the predicted CRMCs in both genomes achieve very high sensitivity for recalling CRM-related elements determined by highly reliable methods, such as the VISTA enhancers and FANTOM promoters. Eighth, recalling CRMs in both genomes are about twice as long as the recalled VISTA enhancers, and the unshared positions in the recalling CRMs are subject to strong evolutionary constrains, while unshared positions in the recalled VISTA enhancers are not. Therefore, most of the predicted CRMCs in both genomes are likely in correct full-lengths, particularly, those with higher *S_CRM_* scores and lower p-values, while some VISTA enhancers might be only components of long CRMs, but still are at least partially functional (Davidson 2006; Hnisz et al. 2013). However, a small portion of the predicted CRMCs in both genomes might be short components of long CRMs, particularly, those with low *S_CRM_* scores and higher p-values. Clearly, more TF ChIP-seq data are needed to cover the relevant genome regions to predict them in full-lengths.

Nineth, the predicted CRMCs in both genomes are substantially more complete and more accurate than those predicted by other state-of-the-art methods measured by evolutionary constraints (Figure 7C) and sensitivity for recalling experimentally determined VISTA enhancers and FANTOM5 promoters. Thus, dePCRM2 is a powerful and robust method for *de novo* prediction of CRMs and TFBSs in large mammal genomes by integrating a very large number of TF ChIP-seq datasets. Finally, although the functional states (TF binding or non-TF-binding) of some CRMs in a cell/tissue type can be predicted based on the overlaps of the CRMCs and TF binding peaks available in the cell type (Ni and Su 2021), functional states of most of the predicted CRMCs in most cell types in both organisms are currently agnostic due to the limited availability of TF ChIP-seq data in most cell types. Fortunately, it has been shown that when the locus of a CRM is accurately anchored by the bindings of key TFs, few epigenetic marks can be an accurate predictor of the functional state of the CRM (Podsiadlo et al. 2013; Kwasnieski et al. 2014; Dogan et al. 2015; Kleftogiannis et al. 2015; Arbel et al. 2019). Thus, the second step of our proposed two-step approach is to predict the functional states of all the predicted CRMs in any cell type in an organism using a minimal set of epigenetic marks collected from the very cell type.

With the availability in the future of even more TF ChIP-seq datasets for more diverse TFs in more diverse cell/tissue types of humans and mice, as well as of other important model organisms such as *Caenorhabditis elegans, Drosophila melanogaster* and *Arabidopsis thaliana,* we are hopeful to predict even more accurate and complete maps of CRMs and constituent TFBSs in all these genomes. These maps will facilitate characterizing functional states and target genes of the CRMs in various cell/tissue types of the organisms, and elucidating the rules of organization and evolution of CRMs and constituent TFBSs at a genome scale.

## MATERIALS AND METHODS

### Datasets

We downloaded the called binding peaks in 9,060 mouse TF ChIP-seq datasets from the Cistrome database (Mei et al. 2017) and filtered out the low-quality peaks with a read depth score less than 20. The datasets are summarized in Table S1. For each called binding peak, we extracted a 1,000bp peak centered on the middle of the peak. To validate our predictions, we downloaded 620 mouse enhancers from the VISTA Enhancer database (Visel et al. 2007), a total of 49,385 mouse enhancers and 163,311 mouse promoters from the FANTOM5 data portal (Forrest et al. 2014; Lizio et al. 2019), and 2,208 QTLs from Mouse Genome Informatics (MGI) databases (Bult et al. 2019). Two compared our predictions with existing methods, we downloaded 339,815 mouse cCREs (Moore et al. 2020) and 519,386 mouse EnhancerAtlas enhancers (Gao and Qian 2020) from the respective websites.

### Measurement of the overlap of binding peaks between two different datasets

We calculate an overlap score *S*_0_(*d_i_*, *d_j_*) of binding peaks between each pair of datasets *d_i_* and *d_j_*, defined as,

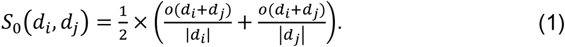

### Prediction of CRMs and constituent TFBSs

To predict CRMs and constituent TFBSs in the mouse genome, we applied the dePCRM2 pipeline (Ni and Su 2021) to the datasets containing 1,000bp peaks. Briefly, we first identify motifs using ProSampler (Li et al. 2019). Secondly, we find the highly frequently co-occurring motifs pairs (CPs) in each dataset by computing a co-occurring score, defined as

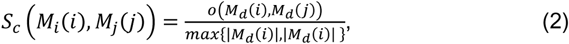

where |*M_d_*(*i*)| and |*M_d_* (*j*)| are the number of binding peaks containing TFBSs of motifs *M_d_*(*i*) and *M_d_* (*j*), respectively; and *o*(*M_d_*(*i*), *M_d_*(*j*)) the number of binding peaks containing TFBSs of both the motifs in *d*. Thirdly, we cluster highly similar motifs in CPs across all the datasets, and find a representative motif in each resulting motif cluster as a unique motif (UM) using ProSampler (Li et al. 2019). Fourthly, we construct an interaction network *N* to model cooccurrence patterns of the UMs and interactions between their cognate TFs. In *N*, the nodes are the UMs that are fully connected, and the edge between UMs *U_i_* and *U_j_* is weighted using an interaction score, defined as,

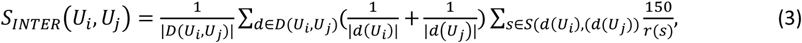

where *D*(*U_i_*, *U_j_*) is the datasets in which TFBSs of both *U_i_* and *U_j_* occur, *d*(*U_k_*) the subset of dataset *d*, containing at least one TFBS of *U_k_*, *S*(*d*(*U_i_*), (*d*(*U_j_*)) the subset of *d* containing TFBSs of both *U_i_* and *U_j_*, and *r*(*s*) the shortest distance between any TFBS of *U_i_* and any TFBS of *U_j_* in a sequence *s* ∈ *S*(*d*(*U_i_*), (*d*(*U_j_*)). Fifthly, we connect two adjacent TFBSs of the UMs if their distance *d ≤* 300bp and predict the connected segment to be a CRM candidate (CRMC) and at the same time, we predict a sequence in the peak-covered regions that cannot be connected to be a non-CRMC. In this way, we partition the peak-covered genome regions in two exclusive sets, i.e., the CRMCs and the non-CRMCs. Sixthly, we evaluate each CRMC containing *n* TFBSs, (*b*_1_, *b*_2_ …, *b_n_*), by computing a CRM score, defined as,

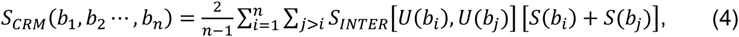

where *U*(*b_k_*) is the UM of TFBS *b_k_*, *S_INTER_*[*U*(*b_i_*), *U*(*b_j_*)] the interaction score between *U*(*b_i_*) and *U*(*b_j_*) in *N*, *S*(*b_k_*) the binding score of *b_k_* based on the position weight matrix (PWM) of *U*(*b_k_*) (Stormo and Fields 1998). Only TFBSs with a positive score are considered. Seventh, we evaluate the statistical significance of each predicted CRMC. To do so, we first generate a Null CRMC set with matched lengths and nucleotide frequencies of the CRMCs using a third order Markov chain model (Li et al. 2019), and a random interaction network *N*′ generated by randomly shuffling the weights in *N*. Then, we compute the *S_CRM_* score for each Null CRMC using formula (4). We compute an empirical p-value for a CRMC with a *S_CRM_*=s, defined as,

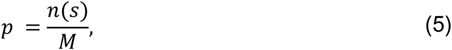

where *n*(*s*) is the number of Null CRMCs with a *S_CRM_* > *s*, and *M* the total number of the CRMCs. Finally, dePCRM predicts functional states (TF-binding or non-TF-binding) in a cell/tissue type of the CRMs whose constituent TFBSs overlap binding peaks of ChIP-ed TFs in the cell/tissue type (Ni and Su 2021).

## AUTHORS’ CONTRIBUTIONS

PN and ZS conceived the project, developed the algorithm, analyzed the data and wrote the manuscript. PN carried out all computational experiments and analysis. PN and DW developed the database. All authors read and approved the final manuscript.

